# Creating an energy efficient central metabolism for boosting biosynthesis without compromising cell growth of yeast

**DOI:** 10.1101/2025.11.22.689960

**Authors:** Xin Ni, Haoyu Wang, Mengyao Zhang, Ning Gao, Jiaoqi Gao, Fan Yang, Yongjin J. Zhou

## Abstract

Long-term natural evolution selects glycolysis as the major metabolic mode for rapid cell growth, which however lacks sufficient NADPH supply to dive the biosynthesis of reduced chemicals such as free fatty acids (FFAs). Engineering energy economical pathway for chemical overproduction always compromises cellular fitness due to the rigidity of cellular metabolism. Here, we successfully replaced glycolysis metabolism with an optimized pentose phosphate pathway (PPP) in an industrial yeast *Ogataea polymorpha*, for the first time, which enabled a higher energy generation efficiency than glycolysis and a balanced supply of ATP and NADPH. More importantly, we discovered a global carbon metabolism regulator *CMR* that drives metabolic flux toward glycolysis for energy generation, and its disruption relieved the tight regulation of metabolic flux distribution and significantly enhanced the efficiency of cellular energy generation, which significantly boosted the FFA production by 63% in a FFA overproducing chassis. The final engineered yeast produced FFAs at a titer of 41.7 g/L, the highest titer reported by microbial fermentation. Our work provides valuable insights into the metabolic regulation mechanisms and a feasible approach for constructing energy efficient metabolism for chemical overproduction.

## Main

Cell growth is fundamentally dependent on a well-balanced flow of carbon through various interconnected metabolic pathways. This process, referred to as carbon budgeting, ensures that cells can generate adequate macromolecules and energy, such as adenosine triphosphate (ATP), to maintain their growth^1^. Natural evolution has led to almost all organisms relying on the Embden–Meyerhof–Parnas pathway (EMP, glycolysis) to generate energy and macromolecules for survival ^2, 3^. This may be because glycolysis is not sensitive to environmental perturbations and changes in cellular demand ^4^.Cellular adaptations that enable the efficient utilization of resources ^5, 6^, and promote the resistance of metabolic pathways to flux alterations and/or genetic modifications, leading to the overproduction of target metabolites ^7^.

There is no evidence that suggests that glycolysis is the optimal pathway for the industrial production of target molecules. For example, excessive glycolysis in yeast often leads to the accumulation of ethanol (Fig. 1)^8, 9^, which makes the production of desired metabolites difficult as a result of imbalanced metabolic flux^5, 10, 11^; thus, exploring alternative metabolic pathways with balanced metabolic flux and energy transfer is necessary.

**Fig. 1:**
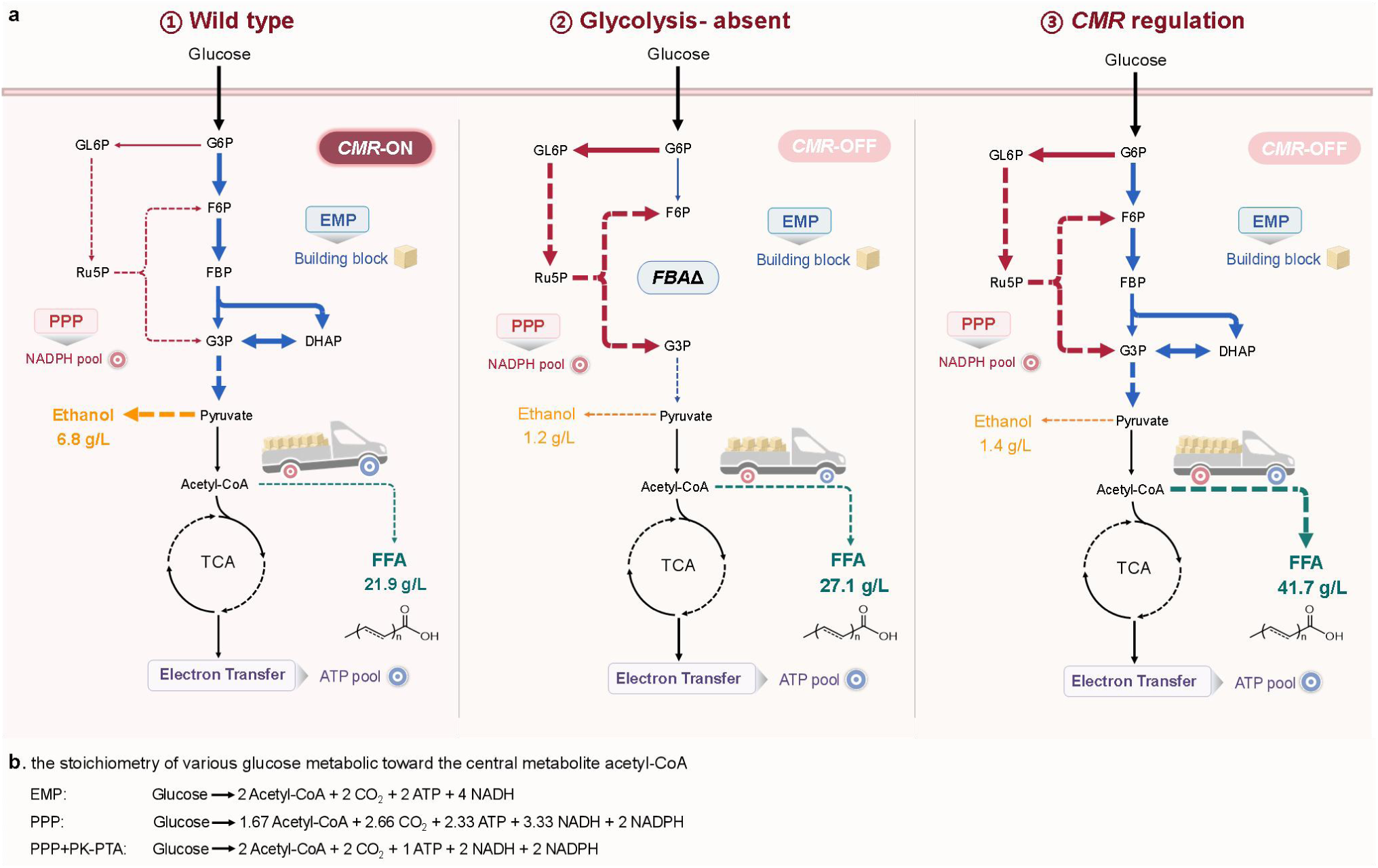
Reshaping the central metabolic network for FFA production by laboratory adaptive evolution and metabolic regulation. (a) The scheme of engineering central metabolism for improved FFA biosynthesis. ① In wild-type *O. polymorpha*, the central carbon metabolism network dominated by glycolysis usually makes redirecting metabolic fluxes toward desired metabolites challenging, resulting in overflow metabolism. ② By blocking glycolysis and conducting laboratory adaptive evolution, new central carbon metabolism is dominated by the pentose phosphate pathway (PPP), which significantly reduces overflow metabolism and increases the yield of FFAs. ③ The key regulatory factor *CMR* has been identified, and the new central metabolic network regulated by *CMR* supplies more balanced NADPH, ATP, and the precursor acetyl-CoA for cell growth and FFA biosynthesis by fine-tuning glycolysis, the PPP, the tri-carboxylic acid (TCA) cycle, and the electron transport chain (ETC). G6P, glucose-6-phosphate; GL6P, gluconate-6-phophsate; F6P, fructosec-6-phosphate; FBP, fructose-1,6-bisphosphate; G3P, glycerol-3-phosphate; DHAP, dihydroxyacetone phosphate. (b) The stoichiometry of various glucose metabolic pathways toward the central metabolite acetyl-CoA.

Most organisms possess alternative metabolic pathways to glycolysis, such as the phosphoketolase pathway^12^, the Entner–Doudoroff (ED) pathway^13^ and the pentose phosphate pathway (PPP) ^14, 15^. Among all alternative glycolytic routes, the PPP plays crucial roles in providing essential cofactors and metabolites for cell growth, such as NADPH as a reducing factor to drive biosynthesis^16, 17^ and 5-phosphate ribose (R5P) as a precursor for nucleotides and coenzymes^18, 19^. Furthermore, the PPP plays a core role in maintaining cellular redox homeostasis, especially during the oxidative stress response ^20, 21^. The PPP branches off from the EMP at glucose-6-phosphate (G6P) and through a series of reactions produces glycerol-3- phosphate (G3P), which can reenter the EMP ^22^ (Fig. 1a). With glucose as a substrate and acetyl-CoA as a product, the EMP produces two ATP molecules, four NADH molecules and two acetyl-CoA molecules per glucose molecule without NADPH generation ^23, 24^, whereas the PPP produces 2.33 ATP molecules, 3.33 NADH molecules and 1.67 acetyl-CoA molecules per glucose with the generation of two NADPH molecules ^25^ (Fig. 1b). Although the EMP produces more energy and precursors per glucose molecule than does the PPP and has lower carbon losses, two NADPH molecules are simultaneously produced by the PPP, which are essential for the biosynthesis of highly reducing molecules such as free fatty acids (FFAs) ^16, 17^. Increasing evidence suggests that increasing the PPP flux improves the microbial biosynthetic capacity ^26–29^. Therefore, we are curious whether the PPP can completely replace the functions of the EMP without compromising cellular fitness, which is highly useful for providing insights into the regulation of cellular central metabolism, as well as the construction of efficient cell factories for sustainable production.

Here, we constructed the sole PPP-growing yeast *O. polymorpha* by blocking the EMP pathway and performing laboratory adaptive evolution (Fig. 1), which increased biomass and FFA production by 29% and 83%, respectively. Genome sequencing and transcriptome characterization revealed that the central metabolism regulator *CMR* (28920315) plays an essential role in regulating the metabolic flux between the EMP and PPP as well as the rearrangement of flux throughout central carbon metabolism, and its inactivation restores the growth of EMP-blocking cells. Deleting *CMR* reshaped the central carbon metabolism network with enhanced PPP flux and improved the efficiency of energy (ATP) generation and protein synthesis, which resulted in the highest microbial production of FFAs at a titer of 41.7 g/L combined with expressing a phosphoketolase pathway (Fig. 1). The results of this study show that cellular metabolism can be rewired for cofactor and efficient biosynthesis by regulating metabolism via the manipulation of a single regulatory factor and provide a feasible approach for constructing efficient cell factories.

## RESULTS

### Growth deficiency of the glycolysis disrupting strain (*fba*Δ) in glucose minimal medium

Yeast naturally metabolizes glucose through the EMP and PPP (Fig. 2a, b), with the EMP taking the major metabolic flux ^24^. Here, we attempted to create a yeast that can metabolize glucose solely through the PPP by blocking the EMP in the potential industrial yeast *O. polymorpha* ^30^, which has a broad substrate utilization capacity ^31^. We used the FFA-producing strain NX1 as a starting strain ^32^ for investigating the metabolic regulation between cell growth and product biosynthesis since the high dependence of NADPH on FFA biosynthesis can respond to PPP regulation ^5, 30, 33^.

**Fig. 2:**
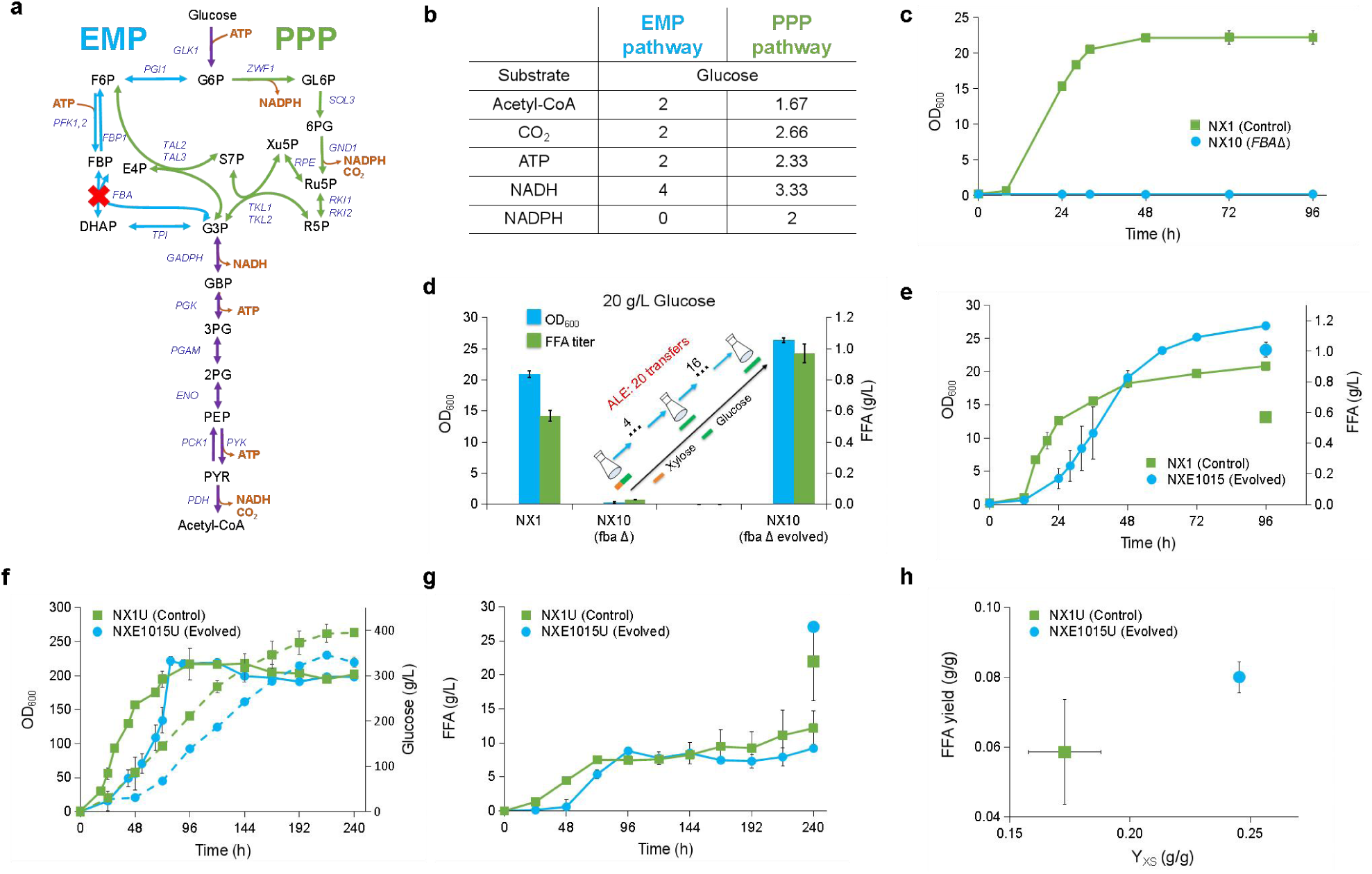
Laboratory adaptive evolution promotes FFA production in glycolytic-deficient strains. (a) Scheme of the EMP and PPP. EMP glycolysis (blue) and PPP glycolysis (green) are parallel, and they share the steps of lower glycolysis (purple). (b) Comparison of carbon exchange and bioenergetics between the EMP and PPP. (c) Cell growth of strain NX10 (*fba*Δ) and the control strain NX1 in G20 minimal medium. (d) Cell growth and FFA production in the control strain, *fba*Δ strain and evolved strain in G20 minimal medium. The *fba*Δ strain had impaired growth in G20 minimal medium, whereas the control strain grew normally (OD_600_= 20). After 20 transfers of ALE, as shown in Figure S1, the growth of the evolved strains was restored, and high-level production of FFAs was detected. (e) Cell growth and FFA production in G20 minimal medium of strain NXE1015 (evolved) and the control strain NX1. (f) Cell growth (solid line) and the consumption of glucose (dashed line) by strains NXE1015 and NX1 during fed-batch fermentation in bioreactors. (g) FFA production of strains NXE1015 and NX1 during fed-batch fermentation in bioreactors. (h) The yield of FFAs and biomass (Y_X/S_) of strains NXE1015 and NX1. The data are presented as the means ± s.e.ms. (n = 3 biologically independent samples for all the experiments except for the fed-batch fermentation, which included 2 biologically independent samples).

Aldolase (*FBA*) plays an essential role in glycolysis ^34, 35^, and a single encoding gene exists in *O. polymorpha* (Fig. 2a), which is thus an ideal target for blocking the EMP. Deletion of *FBA* (strain NX10) resulted in growth deficiency in glucose minimal medium (Fig. 2c), indicating that the natural PPP pathway failed to supply sufficient material and energy for yeast growth. We further evaluated the growth of NX10 cells in xylose medium since xylose metabolism relies more on the PPP than does glycolysis (Extended Data Fig. 1a) ^30^. Interestingly, strain NX10 can grow in both minimal xylose medium (X20) and minimal medium containing xylose and glucose (G10X10) (Extended Data Fig. 1b), even if the lag time is longer than that of the control strain NX1. These results suggest that the PPP can produce the material and energy for yeast growth in the presence of glucose and that glucose cultivation has limited metabolic flux toward the PPP pathway.

### Adaptive laboratory evolution restored the growth of the *fba*Δ strain and increased FFA production

We then attempted to restore the growth of strain NX10 (*fba*Δ) in glucose minimal medium through adaptive laboratory evolution. Five independent NX10 colonies were cultivated for four transfers in minimal medium with 10 g/L glucose and 1 g/L xylose (G10X1) and then cultivated in glucose medium (G10) for 16 transfers (Fig. 2d and Extended Data Fig. 1c, d). All five groups restored cell growth in G10 medium with glucose as the sole carbon source (Extended Data Fig. 1e), with various FFA production levels (Extended Data Fig. 1f). Compared with the control strain NX1, all the selected evolved strains presented a lower specific maximum growth rate (μ^max^) or longer lag time but greater FFA production and final biomass (except NX4-4) (Extended Data Fig. 2). These results indicated that the evolved *fba*Δ strain rewired the metabolic network for FFA biosynthesis. The colony NXE1015, which presented relatively greater final biomass and FFA production (1.04 g/L), was selected for further study (Fig. 2e).

We then compared cell growth and FFA production under fed-batch bioreactor cultivation by using the prototrophic strains NX1U (control) and NXE1015U (evolved *fba*Δ strain), which were constructed by *in situ* complementation of the auxotrophic marker gene *URA3* (Ref. ^36^) in strains NX1 and NXE1015, respectively. Complementing *URA3* significantly improved cell growth and FFA production in shaker flasks, especially for the strain NXE1015U, which can produce 1.43 g/L FFAs in shaker flasks (Extended Data Fig. 3a). However, with a similar final biomass, strain NXE1015U still presented a significantly longer lag time (Fig. 2f) and less glucose consumption (Fig. 2g) than that of the control strain NX1U in the bioreactor. Strain NXE1015U produced 27.0 g/L FFAs, which was 1.23 times greater than that of strain NX1U (Fig. 2h). Furthermore, compared with strain NX1U, the evolved strain NXE1015U significantly increased the FFA yield and biomass yield (Y_XS_) (Fig. 2i).

### Metabolic profiling reveals increased PPP pathway activity and ATP production in the evolved strain

We further examined the metabolic flux of strains NXE1015 and NX1 by using isotope labeling (Fig. 3a). Metabolizing 1-^13^C-labeled glucose through the EMP should result in equal (50%) percentages of the labeled (M+1) and unlabeled (M+0) C_3_ metabolites G3P and 3PG, whereas the PPP should result in 100% M+0 metabolites since ^13^C labeling in the first carbon atom of glucose is released as CO_2_ through the PPP (Fig. 3b). Metabolic analysis revealed that the proportion of M+0 was nearly 100% of all the detected EMP metabolites in strain NXE1015 but lower than 80% in strain NX1 (Fig. 3b), which suggested that glucose was metabolized mainly through the PPP in the evolved strain NXE1015 and that the PPP pathway in the control strain NX1 metabolized 60% of the glucose. Similarly, the proportion of M+0 was nearly 100% of all the detected TCA metabolites in strain NXE1015, which was also much greater than that in strain NX1 (Fig. 3b); more importantly, the NADPH/NADP^+^ ratio in strain NXE1015 was three times greater than that in strain NX1 (Fig. 3c). These results proved that we successfully achieved the transition of the flux from the EMP to the PPP, and we constructed a yeast that relies solely on the natural PPP and has increased biomass yield and FFA production.

**Fig. 3:**
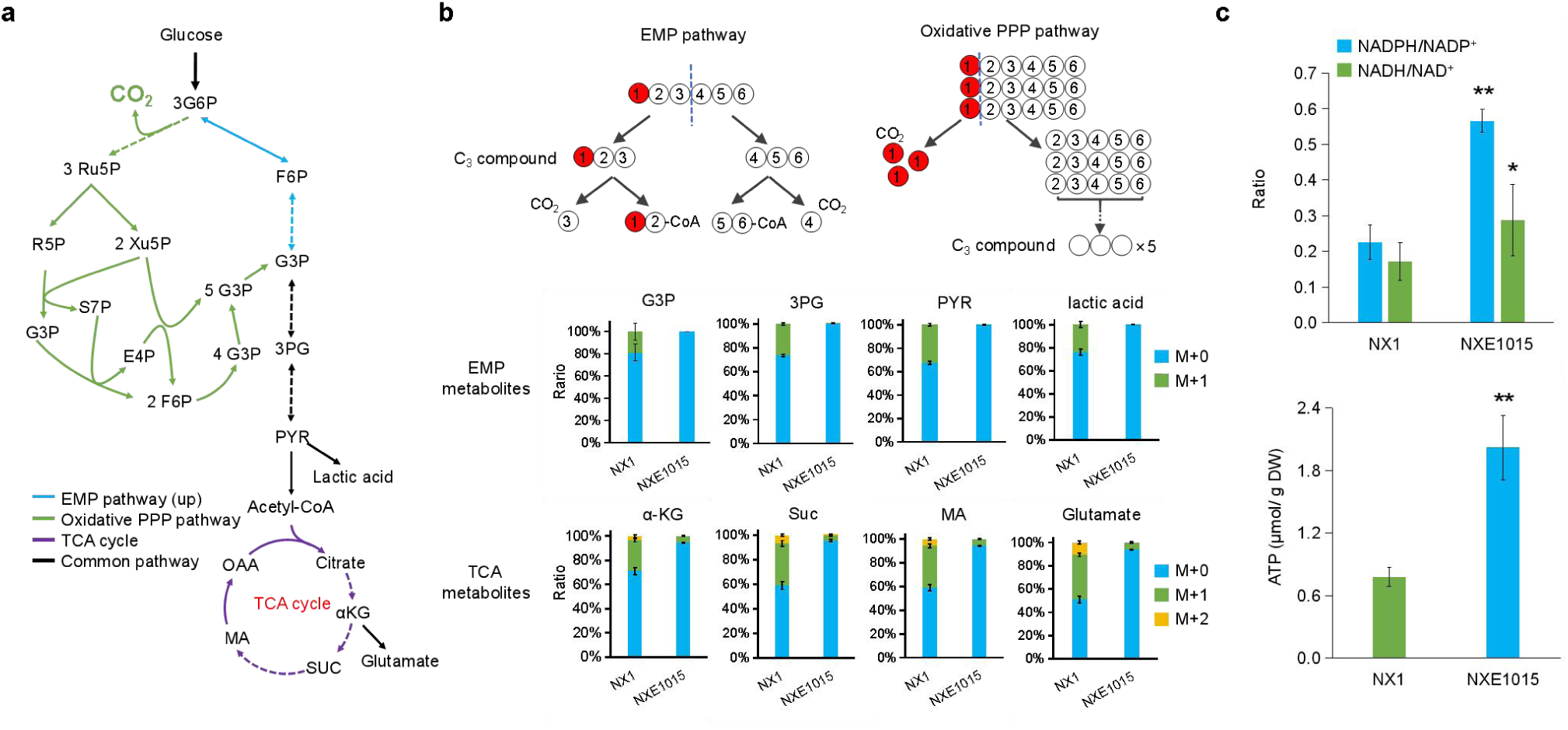
^13^C-labeled glucose tracers reveal the switch from the EMP to PPP. (a) The EMP pathway (blue) and the oxidative PPP pathway (red) are parallel, and they share the steps of lower glycolysis (black) and the TCA cycle (purple). (b) Pathway switching in strain NXE1015 (evolved) was validated by an isotope tracing experiment. One hundred percent 1-^13^C-labeled glucose was used as the carbon source in the G10 minimal medium. The circles indicate the carbon atoms of glucose, and the red dots represent the 1-^13^C-labeled carbon atoms. The blue dashed line represents the breakdown of the carbon–carbon bond. The black dashed line represents multistep pathways. (c) The cofactor measurements of strains NXE1015 (evolved) and NX1 (control), including the NADPH/NADP^+^ ratio, the NADH/NAD^+^ ratio and the ATP level. The data are presented as the means ± s.e.ms. (n = 3 biologically independent samples).

Interestingly, we observed that strain NXE1015 had much higher NADH/NAD^+^ and ATP levels than strain NX1 did (Fig. 3c), which suggested that the reducing power and energy supply were increased with global metabolic rewiring in the evolved strain NXE1015. The PPP has a lower theoretical yield of NADH and ATP than the EMP does (Fig. 1b); therefore, we suspected that the transition of flux from the EMP to the PPP reshaped the entire central carbon metabolism (energy metabolism) of the strain to balance the supply of cofactors, precursor metabolites and energy.

### *A3612* acts as a negative central metabolic regulator for switching the EMP to the PPP

To explore the mechanism of the metabolic shift from the EMP to the PPP, we sequenced at least one independent evolved clone with the best performance from each group by deep sequencing of the wild-type strain as the reference. Notably, single nucleotide polymorphisms (SNPs) and small insertions/deletions (InDels) were identified in the open reading frames, among which the putative transcription factor gene *A3612* (Gene ID: 28920315) was mutated in all five groups (Fig. 4a). Homology alignment searching in the NCBI (https://www.ncbi.nlm.nih.gov/) failed to find any proteins with high sequence similarity to the A3612 protein. Although the transcriptional activator Gal4 from *Saccharomyces cerevisiae* ^37^ had the highest identity (28.8% in 54% of the query cover), the A3612 protein is not a galactose metabolic regulator since *O. polymorpha* cannot metabolize galactose ^38, 39^.

**Fig. 4:**
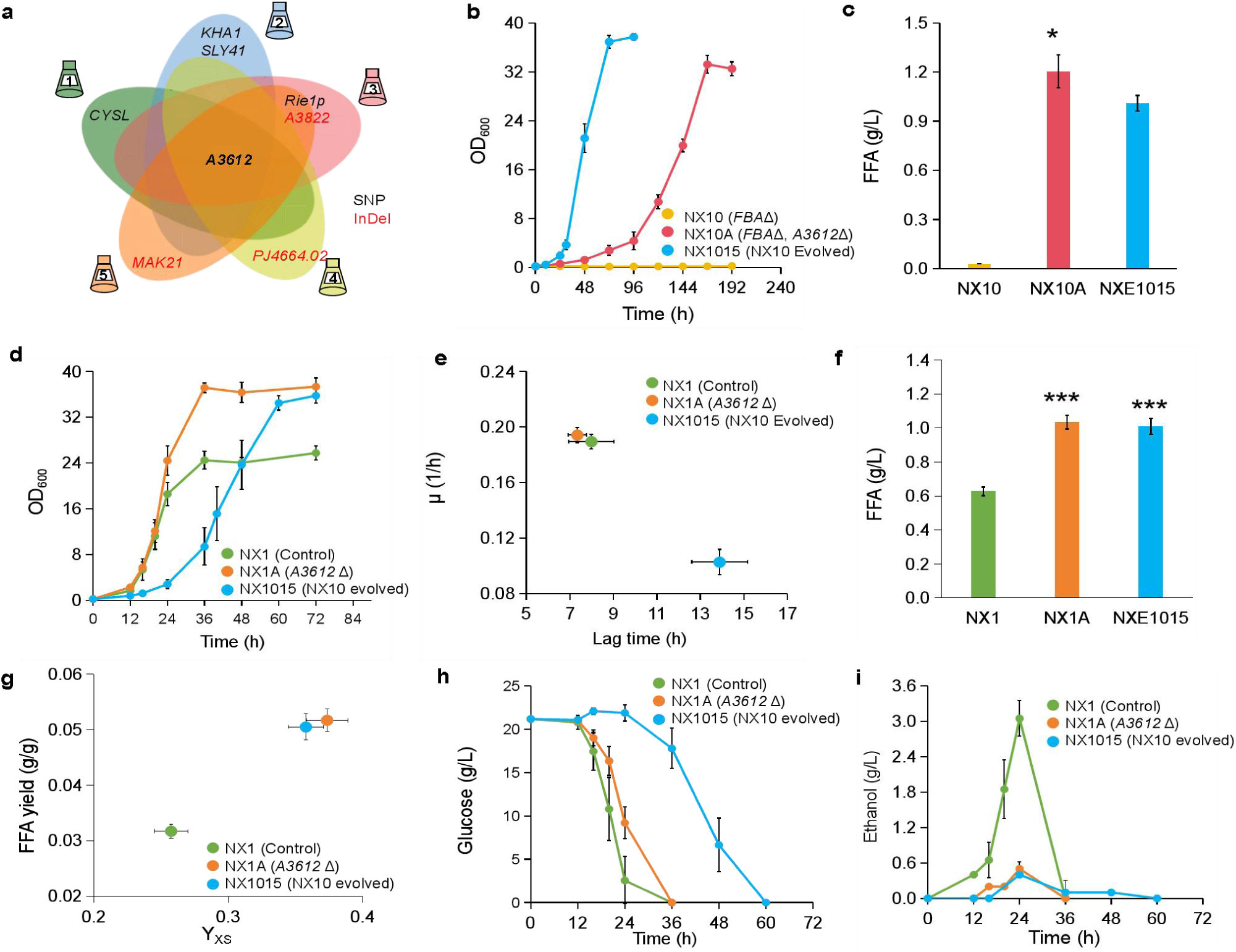
Genome sequencing of evolved strains revealed a key mutation that restored the growth of the *fba*Δ strain and enhanced the growth and FFA production of the strain in G20 medium. (a) Specific mutations (SNPs and InDels) detected via genome sequencing from five groups. Eight mutated genes were identified, among which the putative transcription factor gene *A3612* was shown to be the key mutation. (b) Deletion of *A3612* in the *fba*Δ strain partially restored cell growth in G20 medium. (c) FFA production of strains NX10 (*fba*Δ), NX10A (NX10 *A3612*Δ) and NXE1015 (evolved) in G20 medium. (d) Cell growth of strains NXE1015 (evolved), NX1 (control) and the reverse engineered strain NX1A (NX1 Δ*A3612*). (e) Lag time and growth rate (μ) of strains NX1A and NXE1015 compared with those of strain NX1. (f) FFA production of strains NXE1015, NX1 and NX1A in G20 medium. (g) The FFA and biomass (Y_XS_) yields of strains NXE1015, NX1 and NX1A. (h) The consumption of glucose by strains NXE1015, NX1 and NX1A during fermentation in G20 medium. (i) The accumulation of ethanol in strains NXE1015, NX1 and NX1A during fermentation in G20 medium. The data are presented as the means ± s.e.ms. (n = 3 biologically independent samples).

Deleting *A3612* (strain NX10A) partially restored the growth of the *fba*Δ strain NX10 in glucose medium (Fig. 4b), which implied that this gene was involved in switching the metabolic flux from the EMP to the PPP. Interestingly, with a longer lag time, strain NX10A produced 1.2 g/L FFAs, which was much greater than that of the evolved strain NXE1015 (Fig. 4c). Given the crucial role of glycolysis in cell growth, we attempted to delete the *A3612* gene in strain NX1, which is associated with functional glycolysis. Interestingly, the resulting strain NX1A had a significantly greater final biomass (OD_600_=37) than the control strain NX1 without compromising the growth rate (Fig. 4d, e). In addition, the FFA production of strain NX1A also significantly increased to 1.03 g/L, which was 1.65 times greater than that of strain NX1 and equivalent to that of strain NXE1015 (Fig. 4f). Furthermore, the FFA and biomass yields in strain NX1A were much greater than those in strain NX1 and even slightly greater than those in strain NXE1015 (Fig. 4g). Strain NX1 had the fastest glucose utilization rate (Fig. 4h) and highest accumulation of ethanol during the log phase (Fig. 4i). In contrast, the glucose utilization rate of strain NX1A was slower (Fig. 4h), and that of strain NXE1015 was the slowest, but no obvious ethanol accumulation occurred during the fermentation of these two strains (Fig. 4i). The results indicated that even if *O. polymorpha* is considered a Crabtree negative yeast ^40, 41^, the high glycolytic flux results in overflow metabolism and leads to ethanol accumulation ^42, 43^. *A3612* disruption drives more flux to the PPP, which might be beneficial for FFA biosynthesis and cell growth with relieved overflow metabolism ^44–46^. Interestingly, we also found that the FFA composition in strains NX1A and NXE1015 were similar (Extended Data Fig. 4a), such as linoleic acid, whose proportion was significantly greater than that in strain NX1, which might be related to changes in cofactor levels (Extended Data Fig. 4b) ^38^.

### *A3612* deletion results in significant rewiring of central metabolism

Therefore, to further explore the functions of *A3612*, we conducted whole-transcriptome analysis ^38^ to identify essential differentially expressed genes (DEGs) between the evolved strain NXE1015, the reverse engineered strain NX1A and the control strain NX1 in G20 medium. We found that the number of DEGs in strain NXE1015 was significantly greater than that in strain NX1A (Extended Data Fig. 5a), which might be the result of strain NXE1015 continuously adapting to the environment during long-term evolution. In addition, we found that over 55% of the DEGs were downregulated, especially in strain NX1A (Extended Data Fig. 5a), which may be beneficial for the redistribution of protein resources ^38^. Strains NXE1015 and NX1A presented similar KEGG-annotated DEGs related to metabolic processes (Extended Data Fig. 5b and 5c), especially pathways related to carbohydrate metabolism and amino acid biosynthesis. Notably, the KEGG-annotated DEGs in strain NX1A also included more pathways, such as CoA biosynthesis and amino acid metabolism (Extended Data Fig. 5c), which are believed to be important for the enhancement of cell growth ^47, 48^ and FFA production ^33^.

**Fig. 5:**
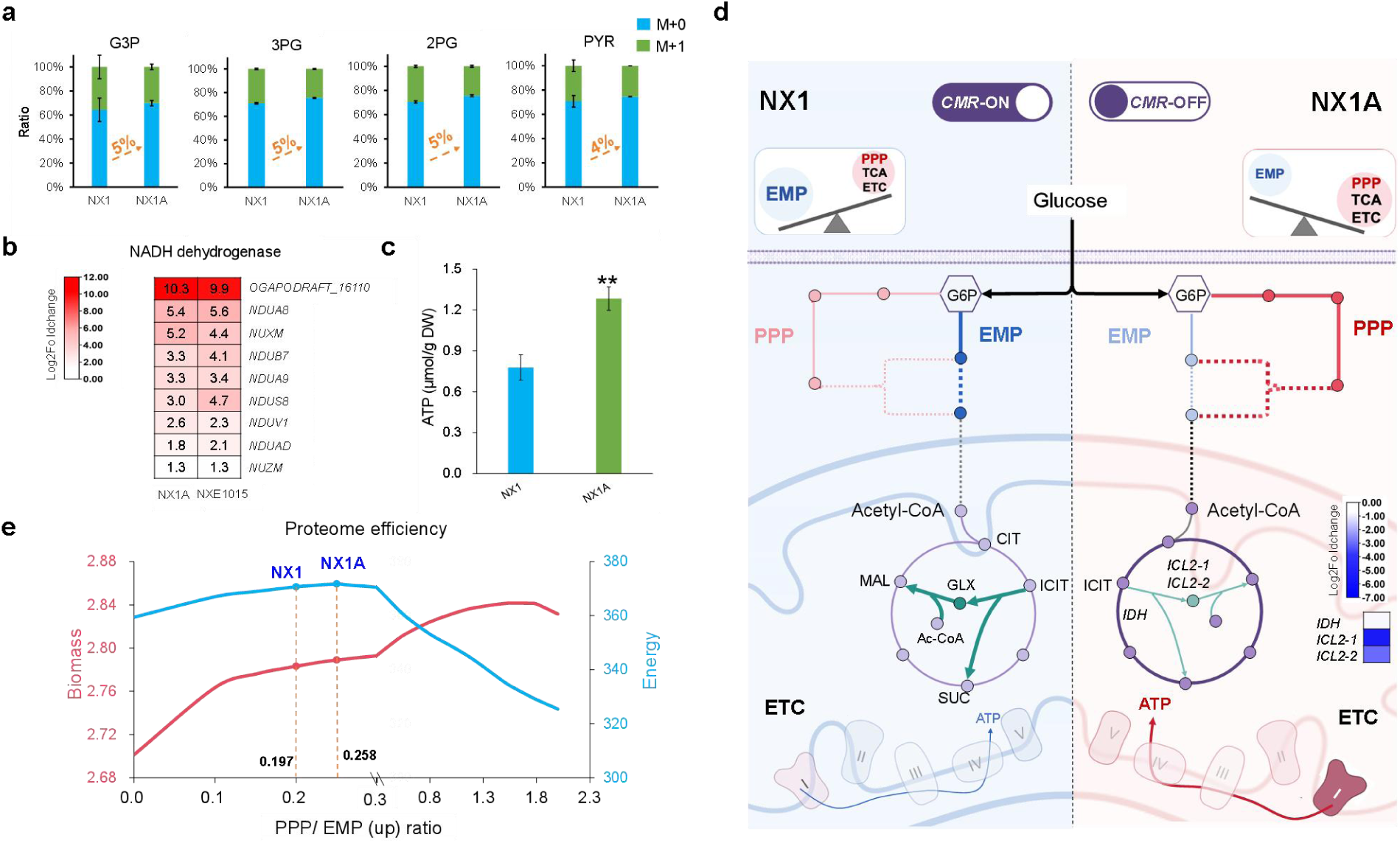
*CMR*, as a switch, can reshape central carbon metabolism. (a) Pathway switching in strain NX1A (reverse engineered strain, Δ*CMR*) was validated by isotope tracing experiments. (b) Transcriptome analysis of the electron transport chain (ETC). Transcriptome data revealed significant upregulation of NADH dehydrogenase in respiratory chain complex I. (c) ATP levels in strains NX1A and NX1. (d) The reshaping of central carbon metabolism regulated by *CMR*. (e) and (f) Protein efficiency with different ratios of PPP and “higher glycolysis” (EMP-up) used for the production of biomass and energy. The data are presented as the means ± s.e.ms. (n = 3 biologically independent samples).

The transcript levels of central carbon metabolism genes in strains NXE1015 and NX1A were surprisingly similar to those in strain NX1 (Extended Data Fig. 6), which further confirmed that the performance of the evolved strain was caused by mutations in gene *A3612*. The EMP genes, such as *PFK1*, *PGAM* and *ENO*, presented decreased transcript levels, which was consistent with the slower rate of glucose utilization in strain NX1A than in strain NX1 (Fig. 4h). We also found that the transcript levels of genes related to ethanol accumulation, such as *PDC*, *ADH2* and *ADH7*, were significantly decreased, which could explain why ethanol accumulation in strains NXE1015 and NX1A was significantly decreased (Fig. 4i). In contrast, several genes were upregulated, including *ACS1* (encoding acetyl-coenzyme A synthetase), *ADH6* (encoding alcohol dehydrogenase) and *YHM2* (encoding citrate carrier protein), which play key roles in acetyl-CoA synthesis and citric acid transport and might be helpful for increasing FFA production by increasing the acetyl-CoA supply ^5, 33^ (Extended Data Fig. 6). We did not find significant changes in the transcription levels of the genes involved in the FFA biosynthetic pathway (Extended Data Fig. 6), indicating that the higher yield of FFA production was caused by the more sufficient precursor acetyl-CoA and reducing power of NADPH rather than the increased FFA synthesis ^33^. On the basis of the obvious regulatory effect of *A3612* on the central carbon metabolism network, we renamed the transcription factor *A3612* carbon metabolism regulator (*CMR*).

### Central carbon metabolism reshaping improves cellular performance via proteome resource optimization

Although ^13^C metabolic analysis revealed that the evolved glycolysis disruption strain NX1015 redirected the flux to the PPP from the EMP, the transcription levels of genes in both the EMP and PPP, particularly the entry genes *PGI1* and *ZWF1* of these two pathways, were downregulated (Extended Data Fig. 6). Therefore, we further examined the flux of the strains NX1A and NX1 by using isotope labeling. The results revealed that the proportion of M+0 among all the detected EMP metabolites in strain NX1A increased by 5% compared with that in strain NX1 (Fig. 5a), indicating that the metabolic flux of the PPP was greater in strain NX1A. In addition, flux balance analysis (FBA) revealed that, compared with that of the control strain NX1, the flux of the PPP increased nearly two-fold in strains NXE1015 and NX1A, with decreased EMP flux (Extended Data Fig. 7). These results proved that the *CMR* plays an important role in regulating the fluxes of the EMP and PPP and that *CMR* deletion drives more flux to the PPP from the EMP.

We then examined the effect of *CMR* deletion on energy metabolism. We found that the transcription levels of complex I genes, which are mainly responsible for NADH dehydrogenation in the respiratory chain ^49^, were significantly increased (Fig. 5b). These upregulated genes suggested that the NADH supply in strains NX1A and NXE1015 was greater than that in the other strains. Consistently, a greater NADH supply resulted in higher ATP levels in strains NX1A and NXE1015 than in strain NX1 (Fig. 3c and 5c). As important nodes for the production of NADH, isocitric dehydrogenase (IDH) and isocitric lyase (ICL) determine the destination of isocitric acid ^50^. Although the transcript levels of *IDH1* and *ICL1/2* both decreased, the decrease in the latter was more significant (Fig. 5d), which led to more isocitric acid entering the TCA cycle rather than the glyoxylate cycle ^51^, thereby increasing the production of NADH for ATP production.

Omics analyses revealed that *CMR* disruption globally reshaped central carbon metabolism by modulating the flux ratio between the PPP and EMP rather than affecting a single metabolic node. These findings prompted investigations into whether a more systematic mechanism underlies the enhanced performance observed in *CMR*-disrupted strains (NXE1015 and NX1A). Recent studies ^52–54^ have demonstrated that proteome allocation plays an important role in determining complex cellular phenotypes. Therefore, we employed an enzyme-constrained metabolic model of *O. polymorpha* ^55^ to investigate the relationship between proteome efficiency and the PPP/EMP flux ratio. We calculated the theoretical maximum proteome efficiency for two distinct cellular objectives: maximal ATP generation and maximal growth. We found that the theoretical maximum proteome efficiency for maximal biomass accumulation initially increased with an increasing PPP/EMP ratio, reaching an optimum value between 1.8 and 2.0, before rapidly declining (Fig. 5e). Although a higher PPP/EMP ratio enhances the supply of precursors and cofactors for cell growth, an excessively high ratio of PPP/EMP may cause redox imbalance. Conversely, the theoretical maximum proteome efficiency for maximal energy supply reached the highest level at a PPP/EMP ratio of 0.3 (Fig. 5e), suggesting that a sufficient EMP is essential for generating NADH and ATP ^44, 56^. Interestingly, the evolved strain NXE1015, with a PPP/EMP ratio of 0.258 (Fig. 5f), had greater theoretical maximum proteome efficiency for both cell growth and energy generation than did the control strain NX1 (PPP/EMP=0.197). This improvement is likely a consequence of reshaped central carbon metabolism. As glycolysis decreases, a large amount of protein resources consumed by glycolysis ^44, 56^ can be redistributed to the PPP and respiration (Fig. 5d). Compared with glycolysis, the PPP can produce more precursors and cofactors, whereas respiration uses a strategy of maximizing energy production to produce ATP ^44^, thus further promoting a sufficient supply of precursors, cofactors, and energy. Consequently, strain NX1A can allocate protein resources more efficiently to generate biomass and energy, thereby demonstrating enhanced performance in both growth and FFA production.

### Enhancing acetyl-CoA biosynthesis increases FFA production

*CMR* deletion drives metabolic flux toward the PPP by providing sufficient NADPH for FFA biosynthesis, which results in an imbalance in the supply of the precursor acetyl-CoA (Fig. 6a). We thus reconstructed a phosphoketolase**–**phosphotransacetylase (PK**–**PTA) pathway ^57, 58^ to divert carbon flux from the PPP by acetyl phosphate (AcP) by expressing *BbPK* from *Bifidobacterium breve* and *CkPTA* from *Clostridium kluyveri* (Fig. 6b). Compared with strain NX1A (Figure 6C), the resulting strain XN8 produced 20% more FFAs (1.21 g/L), with similar FFA profiles (Extended Data Fig. 8). Interestingly, the NADPH/NADP^+^ ratio of strain XN8 was significantly greater than that of strain NX1A (Fig. 6d), and isotope labeling revealed that the M+0 ratio of all the detected substances was greater in strain XN8 than in strain NX1A (Extended Data Fig. 9), suggesting that the expression of *PK***–***PTA* further pulled flux toward the PPP. In addition, the expression of *PK***–***PTA* did not affect the ratio of NADH/NAD^+^ (Fig. 6d), indicating that the energy supply was sufficient in the cell even if more precursors were used for FFA production.

**Fig. 6:**
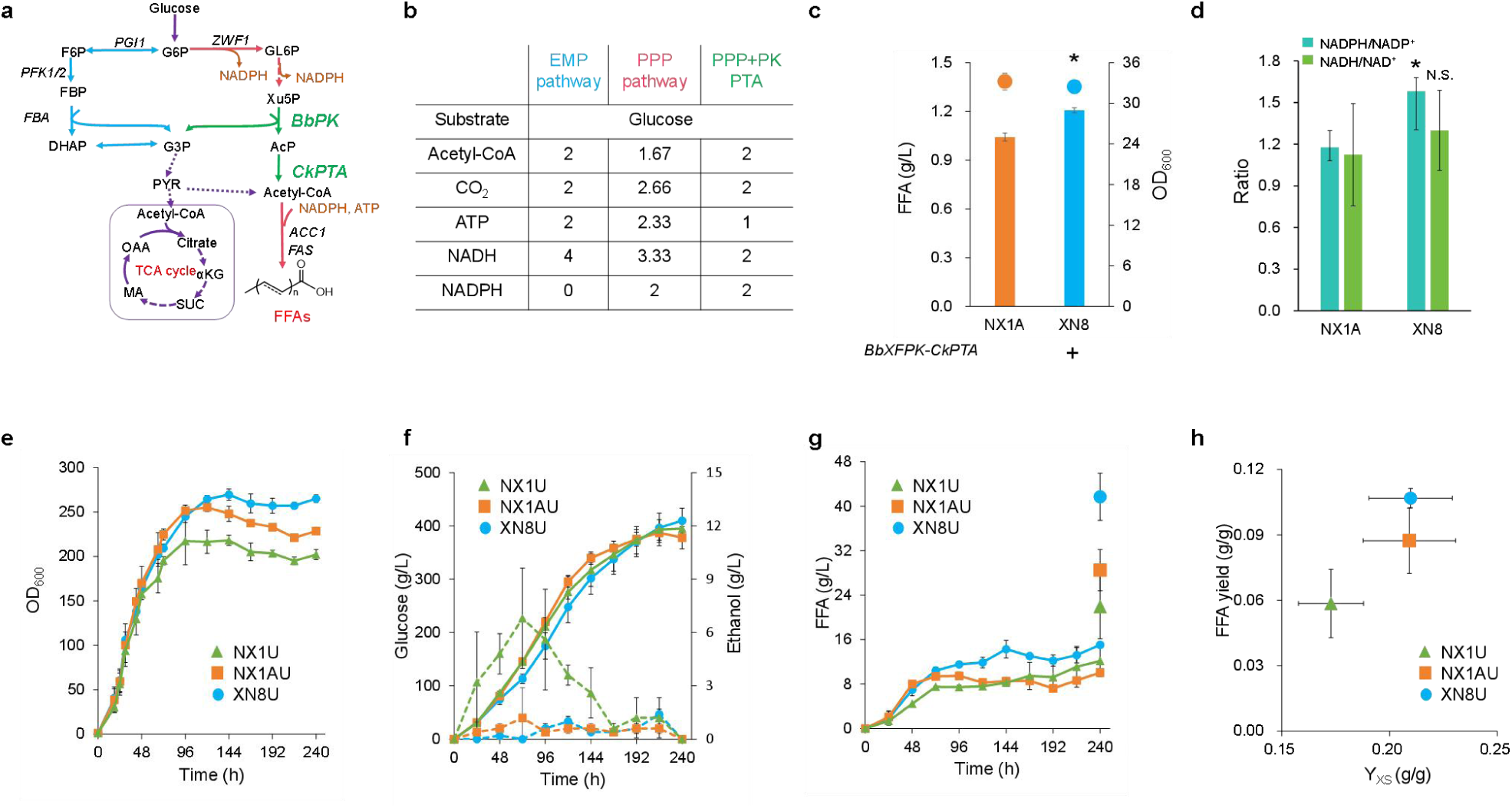
Enhancing acetyl-CoA biosynthesis for improving FFA production. (a) The phosphoketolase–phosphotransacetylase (PK–PTA) pathway can produce acetyl-CoA in the cytoplasm from Xu5P to increase the supply of acetyl-CoA for FFA production. (b) Comparison of carbon exchange and bioenergetics among the EMP, PPP and PPP+PK-PTA. (c) Biomass accumulation and FFA production of strains XN8 (*BbXFPK*-*CkPTA*) and NX1A (control). (d) Cofactor measurements of strains XN8 and NX1A. (e) Cell growth of strains XN8U, NX1AU and NX1U during fed-batch fermentation in a bioreactor. (f) The consumption of glucose (solid line) and the accumulation of ethanol (dashed line) during fed-batch fermentation in the bioreactor. (g) FFA production of strains XN8U, NX1AU and NX1U during fed-batch fermentation in a bioreactor. All the particles were resuspended in culture medium and extracted with ethyl acetate to calculate the final titers of the FFAs. (h) The FFA and biomass (Y_XS_) yields in strains NX1U, NX1AU and XN8U. The data are presented as the means ± s.e.ms. (n = 3 biologically independent samples for all the experiments except for the fed-batch fermentation, which included 2 biologically independent samples).

Finally, we evaluated strain performance in fed-batch fermentation with the prototrophic strains NX1U, NX1AU and XN8U, which were obtained by supplementing the *URA3* gene into the parental strains NX1, NX1A and XN8, respectively. The final biomass (OD_600_) of strains NX1AU and XN8U increased by 15% and 30%, respectively, compared with that of strain NX1U (Fig. 6e). Glucose consumption was similar among the three strains; however, strains NX1AU and XN8U presented much lower ethanol accumulation than did strain NX1U (Fig. 6f), which suggested that strains NX1AU and XN8U alleviated overflow metabolism because the flux redirected toward the PPP from glycolysis. The FFA production of strains NX1AU and XN8U was significantly greater than that of strain NX1U (Fig. 6g), which proved that strains NX1AU and XN8U can produce more FFAs while growing rapidly. Notably, the reverse metabolically engineered strain NX1AU (28.5 g/L) showed better performance than strain NX1U (21.9 g/L) in terms of FFA production (Fig. 6g), which further proves that enhancing the PPP is beneficial for FFA production. Interestingly, strain XN8U produced the highest reported titer of 41.7 g/L FFAs in engineered microbes (Fig. 6g), which was 1.46 and 1.90 times greater than those of strains NX1AU and NX1U, respectively. Similarly, the FFA yield of strain XN8U (0.106 g/g) was 37.7% and 86.0% greater than those of strains NX1U and NX1AU, respectively (Fig. 6h). This fed-batch fermentation once again proved that the rewired central carbon metabolism network is more efficient for FFA production with a balanced supply of cofactors and precursors.

## DISCUSSION

From an evolutionary perspective, cellular metabolic pathways have been optimized to efficiently utilize resources for cell growth, which led to a reliance on glycolysis to generate energy and biomass precursors for microbial survival in the environment. However, this resource-exclusive metabolic network is not always optimal for the biosynthesis of target products. For example, rapid and redundant glycolysis always involves excessive glucose metabolism with the accumulation of ethanol ^8, 9^, which makes the production of desired metabolites challenging due to metabolic flux imbalance ^5^. Therefore, the various pathways of glucose metabolism must be reevaluated to develop more economical biotransformation methods.

Considering the essential role of NADPH in the biosynthesis of reducing chemicals such as FFAs, we constructed a yeast that can grow solely through the PPP pathway for the first time by metabolic engineering and ALE. We further revealed that the causal mutation was the disruption of the negative transcription factor *A3612* through genome sequencing and reverse metabolic engineering. Although *A3612* shares some sequence similarity with *GAL4* ^37^, which encodes a galactose metabolism regulator in *S. cerevisiae*, the lack of galactose in *O. polymorpha* ^38, 39^ suggests that *A3612* might have a different function than *GAL4*. We found that *A3612* deletion redirected the metabolic flux into the PPP rather than into glycolysis, resulting in a rearrangement of the entire central carbon metabolism, including the downregulation of glycolysis, fermentation, and the glyoxylate cycle and the upregulation of complex I, which is responsible for NADH dehydrogenation. This metabolic reconfiguration led to a more sufficient supply of precursors, cofactors, and energy in cells, leading to significant increases in both biomass and FFA production, even in strains with undisrupted glycolysis. Therefore, we renamed *A3612* the central metabolism regulator (*CMR)* to distinguish it from *GAL4* ^37^ in *S. cerevisiae*.

After calculating and comparing the protein efficiency used for energy and biomass production, we believe that a trade-off ^59^ might exist between the metabolism of material and energy and that the transcription factor *CMR* is the key switch in the trade-off. Specifically, cells can utilize glucose through different metabolic routes to generate cofactors, energy and biosynthetic precursors in different proportions according to their needs ^20^, which is beneficial for quickly adapting to different environments. When *CMR* exists, genes in glycolysis are activated, allowing cells to rapidly generate energy at a low protein cost ^44, 60^, which is beneficial for quickly competing for resources. However, when *CMR* is disrupted or deleted, glycolysis is weakened, and more precursors enter the PPP to produce more NADPH and R5P, which promote the resistance of cells, especially in activated neutrophils ^20^. Moreover, more protein resources are being utilized for respiration to maximize ATP yield ^44^.

Finally, we implemented a metabolic design that further increased the supply of acetyl-CoA precursors for FFA production. The obtained strain produced FFAs at a titer of 41.7 g/L, which was the highest reported titer resulting from microbial fermentation, proving that the new central carbon metabolism network is more beneficial for FFA production. Furthermore, since we succeeded in the high production of FFAs, we are therefore confident that the new central carbon metabolism network could also be used for the production of other acetyl-CoA derivatives ^30, 57^.

In summary, we explored the possibility of the PPP as an alternative pathway to the EMP to support microbial growth and chemical production and identified a transcription factor encoded by *CMR* that can regulate the expression of genes in the central carbon metabolism network, such as the EMP, PPP, and respiration genes. This regulatory function involves the rational allocation of the supply of ATP and macromolecules for efficient energy metabolism. Our work provides valuable insights for a deeper understanding of the metabolic regulatory mechanisms of microorganisms.

## SUPPLEMENTAL INFORMATION

Supplemental information can be found online at

## ACKNOWLEDGEMENTS

This work was supported by the National Natural Science Foundation of China (22425807, 22478382), LiaoNing Revitalization Talents Program (XLYC2402041) and DICP innovation grant (DICP I202335 and DMU-2&DICP UN202505).

## AUTHOR CONTRIBUTIONS

X.N., J.G. and Y.J.Z. designed the research; X.N. designed and performed most of the experiments; X.N., J.G. and H.W. analyzed the data; H.W. provided model calculation; X.N., J.G. and N.G. performed the fed-batch fermentation. M.Z. assisted with experimental methods. X.N., J.G., F.Y. and Y.J.Z. wrote the manuscript. J.G., F.Y. and Y.J.Z. supervised the research.

## DECLARATION OF INTERESTS

The authors have patent filings to disclose: X.N., J.G., and Y.J.Z. have one patent (20251333) for protecting part of the work described herein. All other authors declare no competing financial interests.

## METHODS

### Strains and cultivation

All the strains used in this study are listed in Supplementary Table 1. All strains were stored in our laboratory or constructed in this study. For cultivating yeast strains, YPD medium (20 g/L glucose, 20 g/L peptone, and 10 g/L yeast extract) or Delft minimal medium (2.5 g/L (NH_4_)_2_SO_4_, 14.4 g/L KH_2_PO_4_, 0.5 g/L MgSO_4_•7H_2_O, 1 mL/L vitamin solution, and 2 mL/L trace metal solution) containing the specific substrates were used. For strain screening, synthetic dropout (SD) medium with 6.7 g/L yeast nitrogen base without amino acids and 20 g/L glucose was utilized, supplemented with the necessary amino acids. The *URA3* marker was removed on SD+URA+5-fluoroorotic acid (FOA) plates, which consisted of 6.7 g/L yeast nitrogen base without amino acids, 20 g/L glucose, and 2 g/L 5-FOA (Sangon Biotech, Shanghai, China). *Escherichia coli* DH5α was used to construct plasmids, which were cultivated in LB medium (10 g/L tryptone, 10 g/L NaCl, and 5 g/L yeast extract supplemented with 100 mg/L ampicillin). All the strains were cultivated at 37°C and 220 rpm in a shaking incubator (Zhichu Shaker ZQZY-CS8).

For all the fatty acid production experiments, the strains were precultured in YPD medium for 16–18 h and then washed once with minimal medium, after which they were subsequently transferred into minimal medium (containing 20 g/L glucose) with an initial OD_600_ of 0.2. The strains were cultivated at 37°C and 220 rpm for 96 h prior to measurements of the biomass, sugars, and fatty acids. In this study, biomass was represented by the optical density at 600_nm_, which can be converted to DCW (g/L) with a coefficient of 0.2.

### Genetic manipulation by the CRISPR/Cas9 system

All strains were constructed via our previously established CRISPR/Cas9 system in *O. polymorpha* ^61^. The deletion and integration of genes were carried out following the standard procedure with gRNA design, plasmid and donor DNA construction, transformation, colony verification, and selective marker removal ^62^. The resulting plasmids, primers, donor DNA constructs and optimized heterologous genes are listed in Supplementary Tables 2, 3, 4 and 5, respectively. DNA manipulation, such as PCR amplification, enzyme digestion, and ligation, was performed according to standard procedures. Donor DNA and gene expression cassettes were constructed by overlap extension PCR (Supplementary Table 4). For scarless gene deletion, the upstream homologous arm was directly linked with the downstream homologous arm via overlap extension PCR. Purified DNA fragments (500 ng) were transformed together with a specific gRNA plasmid (500 ng). Similarly, for site-specific integration, each DNA fragment, such as the upstream and downstream homologous arms, promoter, gene, and terminator, was prepared to construct the donor DNA by overlap extension PCR. The other steps were the same as those for scarless gene deletion. *O. polymorpha* was transformed by electroporation.

### Adaptive laboratory evolution

The ALE process is illustrated in Extended Data Fig. 1c. Briefly, the parent strain NX10 was spotted on a YPD plate, and five independent colonies were cultivated in YPD medium at 37°C and 220 rpm for 16–20 h. In the first stage of evolution, these five strains were transferred to minimal medium (G10X1) containing a mixture of glucose and xylose at a ratio of 10:1. If the OD_600_ reached 6–8, the evolved strains were transferred to medium (G10) supplemented with 10 g/L glucose as the sole carbon source. If not, the strains were transferred to the same medium with an initial OD_600_ of 0.4. For the second stage of evolution, these five strains were transferred to G10 medium with an initial OD_600_ of 0.4. If the OD_600_ reached 3–6 within 48 h, the evolved strains were transferred to the same medium with a decreased initial OD_600_. The evolution process was finished when the strains were able to grow normally (achieving an OD_600_ of 5–6 within 36 h) in G10 medium. Every five transfers, the strains were preserved and verified by PCR. The final evolved strains were spotted on a YPD plate, and eight colonies from each group were picked to evaluate the performance of cell growth and FFA production in minimal medium (G20) containing 20 g/L glucose as the sole carbon source.

### Genome sequencing analysis

Five strains were selected for genome sequencing, including the evolved strains in group I (5), group II (2), group III (3), group IV (7) and group V (8). Genomic DNA was extracted by the SDS method, detected by agarose gel electrophoresis and quantified by a Qubit^®^ DNA Assay Kit with a Qubit^®^ 3.0 Fluorometer (Invitrogen, USA). A total of 0.2 μg of DNA per sample was used as input material for the DNA library preparations. The sequencing library was generated by the NEBNext^®^ Ultra^TM^ DNA Library Prep Kit for Illumina (NEB, USA) following the manufacturer’s recommendations, and index codes were added to each sample. Briefly, a genomic DNA sample was fragmented by sonication to a size of 350 bp. Then, the DNA fragments were endpolished, A-tailed, and ligated with the full-length adapter for Illumina sequencing, followed by further PCR amplification. After the PCR products were purified by the AMPure XP system (Beckman Coulter, Beverly, USA), the DNA concentration was measured by a Qubit^®^ 3.0 Fluorometer (Invitrogen, USA), and the size distribution of the libraries was analyzed by an Agilent 2100 Bioanalyzer and quantified by real-time PCR (>2 nM).

Clustering of the index-coded samples was performed on a cBot Cluster Generation System by an Illumina PE Cluster Kit (Illumina, USA) according to the manufacturer’s instructions. After cluster generation, the DNA libraries were sequenced on an Illumina platform, and 150 bp paired-end reads were generated. The original fluorescence image files obtained from the Illumina platform were transformed to short reads (raw data) by base calling, and these short reads were recorded in the FASTQ format, which contains sequence information and corresponding sequencing quality information. Valid sequencing data were mapped to the reference genome by Burrows–Wheeler Aligner (BWA) software to obtain the original mapping results stored in BAM format (parameters: mem -t 4 -k 32 -M). The results were subsequently rearranged for duplication by SAMtools (parameter: rmdup) and Picard (http://broadinstitute.github.io/picard/).

### Transcriptome analysis

To conduct the transcriptional analysis, the control strain (NX1), evolved strain (NXE1015) and reverse metabolic engineering strain (NX1A) were cultured in G20 minimal medium at 37°C and 220 rpm until the OD_600_ reached 3–4. The total RNA of each sample was extracted with a RNeasy Mini Kit (QIAGEN) according to the manufacturer’s instructions. RNA with an integrity of more than 6.5 that was detected by a 2130 Bioanalyzer (Agilent Technologies) was used to construct and sequence the library. The complementary DNA libraries were constructed and sequenced by the BGISEQ-500 platform at the Beijing Genomics Institute. To detect differences in gene expression among various samples, single-end technology was used to obtain approximately 40–50 bp reads in a single run. DEGs were analyzed by the DESeq2 R package (v.1.20.0) and are displayed as average values of three clones from each group.

### Isotope labeling

The minimal medium with 10 g/L 100% D-glucose-1-^13^C (Sigma‒Aldrich) ^20, 63^ in a shake flask was used for sample preparation for the isotope labeling experiment. The strains were inoculated at an OD_600_ of 0.2 and harvested in the mid- to late-exponential phase (OD_600_ of approximately 6–8).

Chromatographic separation was performed on a Thermo Fisher Ultimate 3000 UHPLC system with a Waters BEH amide column (2.1 mm * 100 mm, 1.7 μm). The injection volume was 1 μL, and the flow rate was 0.4 mL/min. The mobile phases consisted of water with 15 mM ammonium acetate and 0.2% ammonium hydroxide (phase A) and 90% acetonitrile (phase B). Linear gradient elution was performed with the following program: 0–0.5 min, 90% B; 8 min, 70% B; 8.1–9.5 min, 50% B; and 9.6–12 min, 90% B. The eluents were analyzed on a Thermo Fisher Q Exactive^TM^ Hybrid Quadrupole-Orbitrap^TM^ mass spectrometer (QE) in heated electrospray ionization negative mode. The spray voltage was set to 4000 V. The capillary and probe heater temperatures were both 320°C. The sheath gas flow rate was 35 (Arb, arbitrary unit), and the auxiliary gas flow rate was 10 (Arb). The S-Lens RF level was 50 (Arb). The full scan was performed at a high resolution of 70000 FWHM (m/z=200) in the range of 70–1050 m/z with the AGC target setting at 3*10^6^.

### Cofactor assay

Strains NX1, NXE1015, NX1A and XN8 were used to determine the intracellular NADP(H) and NAD(H) concentrations according to previous reports ^38^ with some modifications. The cells were cultivated in G20 minimal medium and collected at 24 h. To extract NADP(H) and NAD(H), the cells at a total OD_600_ of 5 were collected and resuspended in 300 μL of 0.2 M NaOH (for NADPH and NADH) or 0.2 M HCl (for NADP^+^ and NAD^+^). Then, glass beads were added and vigorously vortexed at 70 Hz for 120 s three times on an ice-cold holder. This process was repeated five times before heating at 55°C for 10 min. The extracts were neutralized by adding 300 μL of 0.2 M HCl (for NADPH and NADH) or 0.2 M NaOH (for NADP^+^ and NAD^+^) and centrifuged at 12,000 × g for 5 min.

For NADPH and NADP^+^, the cycling assay was performed by using a reagent mixture consisting of equal volumes of 1.0 M bicine buffer (pH of 8.0), 30 mM glucose-6-phosphate, 40 mM EDTA (pH of 8.0), 4.2 mM 3-(4,5-dimethyl-2-thiazolyl)-2,5-diphenyl-2H-tetrazolium bromide, two volumes of 16.6 mM phenazine ethosulfate and three volumes of water. Then, 10 μL of neutralized extract and 180 μL of reagent mixture were added to 96-well plates, and 10 μL of glucose-6-phosphate dehydrogenase (100 U/mL) was added to start the assay. The absorbance at 570 nm was recorded for 10 min (every 30 s for 10 min) at 30°C. For NADH and NAD^+^, the same reagent mixture was used, but 30 mM glucose-6-phosphate was replaced with anhydrous ethanol. Then, 10 μL of neutralized extract and 180 μL of reagent mixture were added to 96-well plates, and 10 μL of alcohol dehydrogenase (500 U/mL) was added to start the assay. The absorbance at 570 nm was recorded for 10 min (every 30 s for 10 min) at 30°C. The slope (ΔA min^−1^) was correlated with the concentrations of cofactors (in mM) by a linear fit equation, which was subsequently used to calculate the concentrations of NADPH, NADP^+^, NADH and NAD^+^ in the extracts from each sample. The total protein concentration was determined by a Nanodrop instrument to normalize the results.

### Determination of ATP

The ATP level was determined by an ATP content assay kit (Solarbio^®^ BC0300) ^64^. The cells were cultivated in G20 minimal medium and collected at 24 h. To extract ATP, the cells at a total OD_600_ of 5 were collected and resuspended in extraction solution. Then, glass beads were added and vigorously vortexed at 70 Hz for 120 s three times on an ice-cold holder. After centrifugation at 10,000 × g for 10 min, 500 μL of chloroform was added to the supernatant, which was subsequently centrifuged at 10,000 × g for 3 min. Then, 100 μL of the supernatant was added to the working solution, and the absorbance A_1_ at 340 nm was recorded immediately. After reacting at 25°C for 3 min, the absorbance A_2_ was measured by using an ATP standard solution with a concentration of 0.625 μmol/mL as the control, the ATP content in the sample was calculated, and the data were normalized to the cell numbers.

### Metabolic flux balance analysis

The genome-scale metabolic model of *O. polymorpha* iUL909 ^65^ was modified by blocking the corresponding reactions to generate strain-specific models for NX1, NX1A and NXE1015. To simulate flux distributions, parsimonious flux balance analysis (pFBA) ^66^ was conducted using these strain-specific models, constrained by experimentally measured glucose uptake, ethanol production, specific growth rates and FFA production rates, with the objective of maximizing ATP maintenance. The calculated fluxes were normalized by dividing each flux by its corresponding glucose uptake rate and multiplying by 100, standardizing the glucose input to 100 units across all strains. Model construction and simulation were performed in Python (v3.9) with COBRApy ^67^ (v0.27).

### Theoretical maximum proteome efficiency

Proteome efficiency is defined as the objective flux per unit of proteome mass (gProt/gDW) ^68^. However, not all proteins present in the cell may be fully utilized for catalytic reactions, implying that some proteins remain inactive or are underutilized. Therefore, the theoretical maximum proteome efficiency (mmol/gProt/gDW) for objective production is defined as the maximum flux per unit of total proteome allocation:

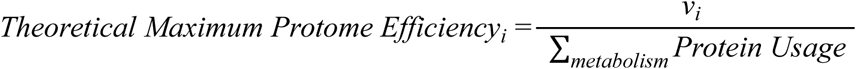

where *v_i_* is the flux of the objective and I,*_metabolism_ Protein Usage* is the total proteome usage involved in metabolism. To calculate the theoretical maximum proteome efficiency, we employ enzyme-constrained genome-scale metabolic models (ecGEMs). Specifically, the ecGEM of *O. polymorpha* emodel_*Ogataea*_*polymorpha*_Posterior_mean.mat ^55^ was used to perform simulations in minimal medium with glucose as the sole carbon source.

The calculation of the theoretical maximum proteome efficiency is formulated as a two-stage linear programming problem:

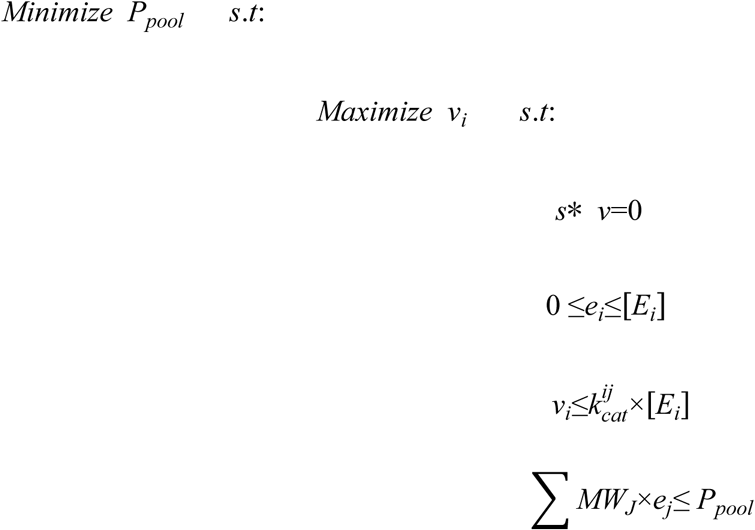

where *P_pool_*represents the total proteome allocation for metabolism, *v_i_* represents ATP generation or biomass reactions, [*E_i_*] represents the enzyme concentration, *e_j_* represents the enzyme usage for reaction J, and *MW_J_* represents the molecular weight of the enzyme involved in reaction J.

Under the condition of minimizing *P_pool_*, all proteins are fully utilized, leading to:

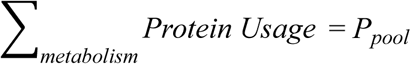

To constrain the flux distribution between the pentose phosphate pathway (PPP) and the Embden–Meyerhof–Parnas pathway (EMP), an additional constraint is introduced:

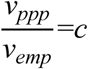

where *v_ppp_*and *v_emp_* are the fluxes through the PPP and EMP, respectively, and is a predetermined constant ratio.

For the simulations, the glucose uptake rate was set as 5 mmol/g DW/h. The constant was varied from 0.2 to 5 to assess the theoretical maximum proteome efficiency across different pathway flux distributions. These simulations were implemented using COBRApy ^67^ (v0.27) in Python (v3.9), with GUROBI (11.0.1) employed as the optimization solver.

### Fed-batch fermentation

Fed-batch fermentation was performed in a DasGip parallel bioreactor system (Eppendorf). The initial batch fermentation was carried out in G20 minimal medium in 1 L bioreactors with a 0.4 L working volume. Precultured engineered strains (OD_600_=5∼6) were inoculated with an initial OD_600_ of approximately 1.0. The temperature, pH and dissolved oxygen (DO) content were set to 37°C, 5.6, and 30%, respectively. The initial agitation speed was set to 400 rpm and increased to a maximum of 800 rpm, depending on the DO level. Aeration was initially provided at 18 sL/h and increased to a maximum of 48 sL/h depending on the DO level. Fed-batch cultivation was started immediately after the carbon source in the batch medium was exhausted, and 5×Delft medium (12.5 g/L (NH_4_)_2_SO_4_, 72 g/L KH_2_PO_4_, 2.5 g/L MgSO_4_•7H_2_O, 10 mL/L trace metals, and 5 mL/L vitamin solution) containing 500 g/L glucose was pumped into the vessels on the basis of a dissolved oxygen tension (DOT) stat control. The residual sugars were monitored to control the feeding rates to maintain the sugar concentration at a low level (less than 15 g/L). During fermentation, fermentation broth was sampled in time to detect residual glycose and ethanol by using a biosensor. After fermentation, all solid FFAs were resuspended in the culture mixture to accurately measure the total FFA titer via ethyl acetate extraction.

### Qualitative and quantitative analysis of fatty acids

Total free fatty acid extraction was modified from previous reports. Briefly, properly diluted cell cultures (100 μL) were mixed with 100 μL ddH_2_O, and then 10 μL of 40% tetrabutylammonium hydroxide (Sigma, Cat No. 86854) was added. Immediately, 200 μL of 200 mM iodomethane (Sigma, Cat No. 18507) in dichloromethane (Sigma) containing 0.1 mg/mL pentadecanoic acid (Sigma, Cat No. P6125) as an internal standard was added, and the mixtures were shaken for 30 min using a vortex mixer (1200 rpm) and then centrifuged to promote phase separation at 2000 × g for 10 min. The dichloromethane layer (150 μL) was transferred into a GC vial with a glass insert and allowed to evaporate to dryness. The extracted methyl esters were then resuspended in pure hexane and analyzed by gas chromatography (Focus GC, Thermo Fisher Scientific) equipped with a Zebron ZB-5MS GUARDIAN capillary column (30 m * 0.25 mm * 0.25 μm, Phenomenex). The GC program was as follows: initial temperature of 40°C, held for 2 min; ramp to 180°C at a rate of 30°C per minute; then, increase to 200°C at a rate of 4°C per min, held for 1 min; and finally, increase to 240°C at a rate of 2°C per minute, held for 10 min. The injection volume was 1 ml. The flow rate of the carrier gas (nitrogen) was set to 1.0 mL/min.

Specifically, for samples of fed-batch bioreactors with visible solids, the whole vessels of cell cultures were extracted with ethyl acetate, the mixtures were shaken at 220 rpm for 30 min, and the upper layer was transferred to a new glass vial and allowed to evaporate to dryness. Then, 200 μL of ddH_2_O was added, and the following procedure was identical to that described above.

### Statistical Analysis

Statistical analysis was performed using Microsoft Excel software using a two-tailed-t-test analysis of variance. Significant differences are marked as *p<0.05, **p<0.01 and ***p<0.001. All the data are presented as the means±s.e.ms. The number of biologically independent samples for each panel was three unless otherwise stated in the figure legends.

---

**Extended Data Fig. 1.**
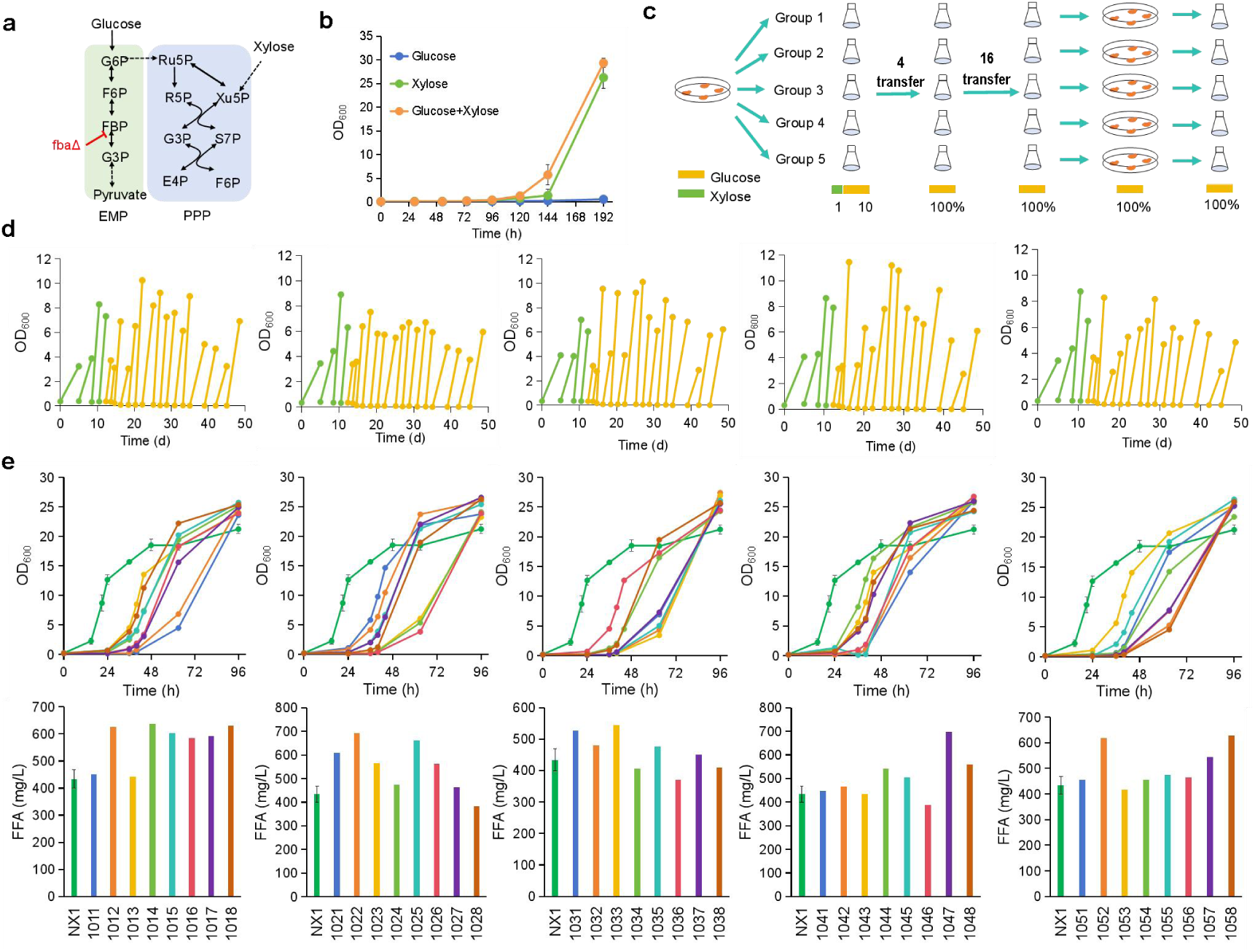
Adaptive laboratory evolution (ALE) of the *fba*Δ strain enabled cell growth and FFA production in glucose. (a) Scheme of the metabolic pathways of glucose and xylose. (b) Cell growth of strain NX10 (*fba*Δ) in minimal medium containing glucose (20 g/L), xylose (20 g/L), or glucose (10 g/L) + xylose (10 g/L). (c) Procedure of ALE. Five independent colonies of strain NX10 were cultivated in G10X1 (10 g/L glucose+1 g/L xylose) medium. If the OD_600_ reached 4∼6 within 48 h, the evolved strains were transferred to G10 medium until they were able to grow normally. The final evolved strains were spotted on YPD plates for further characterization. (d) Procedure of ALE illustrated by cell growth. The final OD_600_ before every transfer was measured, and although distinguished processes had been gone through, all five groups ultimately achieved both cell growth and FFA production in minimal medium containing glucose as the sole carbon source. Eight colonies from each group were selected for cell growth and FFA production (e) in G20 minimal medium. The data are presented as the means ± s.e.ms. (n = 3 biologically independent samples).

**Extended Data Fig. 2.**
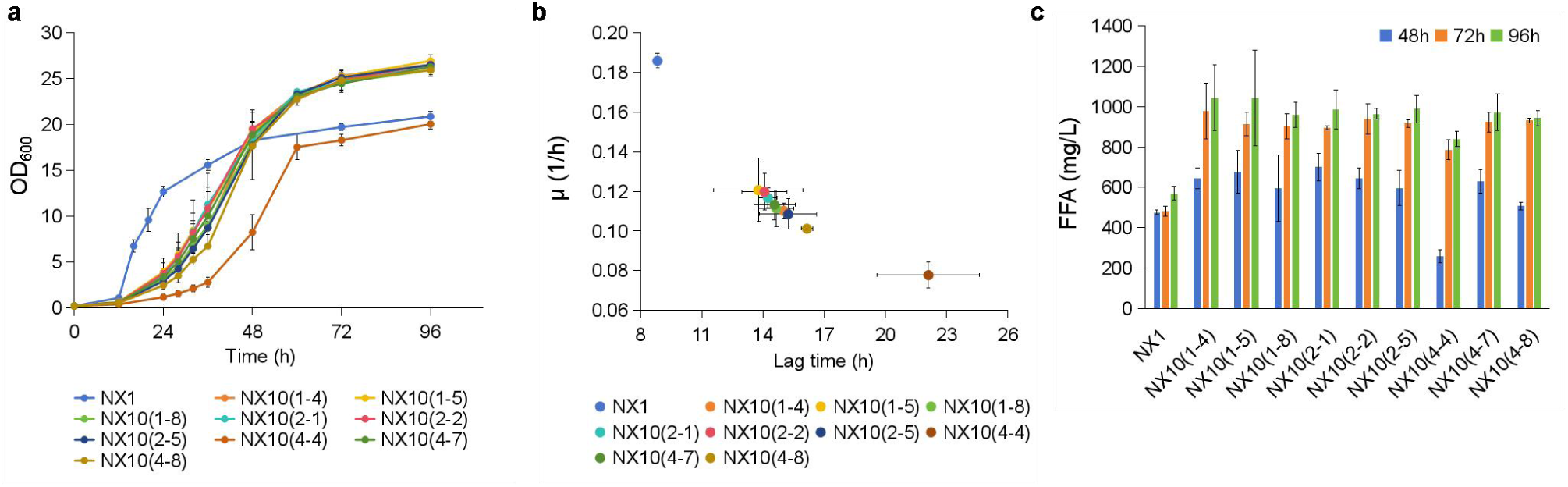
Evaluation of monoclonal strains of evolutionary strains. (a) Cell growth of different evolved strains and strain NX1 in G20 medium. (b) Lag time and growth rate (μ) of different evolved strains and strain NX1 in G20 medium. (c) FFA production of different evolved strains and strain NX1 at different times. The data are presented as the means ± s.e.ms. (n = 3 biologically independent samples).

**Extended Data Fig. 3.**
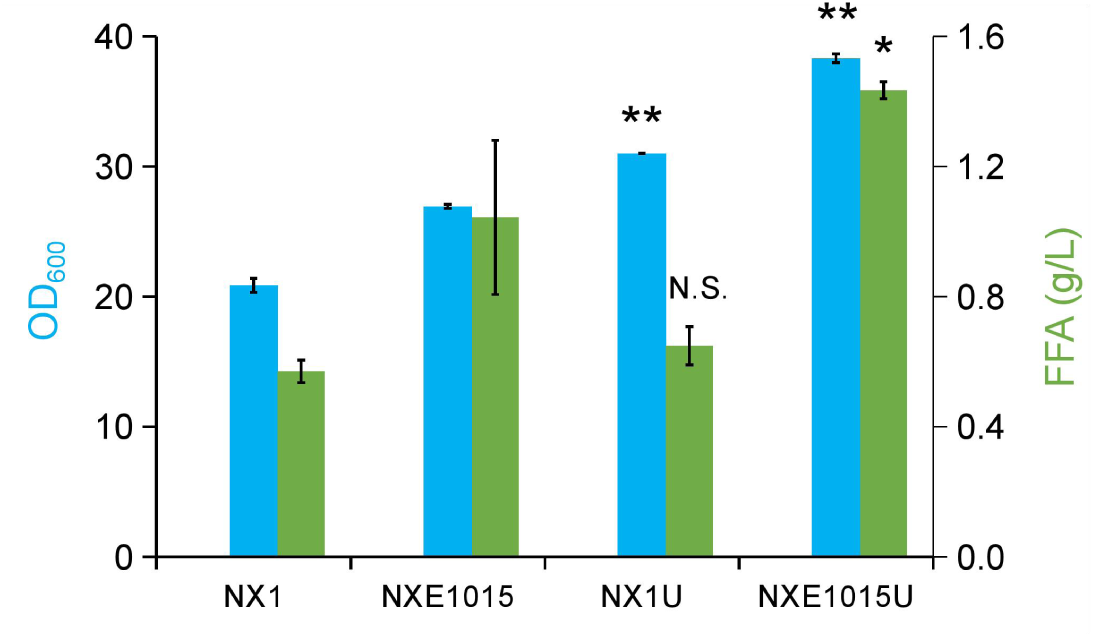
FFA production of the prototrophic strains. The prototrophic strains NX1U (control) and NXE1015U (evolved *fba*Δ strain) were constructed by *in situ* complementation of the auxotrophic marker gene *URA3* in strains NX1 and NXE1015, respectively. The data are presented as the means ± s.e.ms. (n = 3 biologically independent samples).

**Extended Data Fig. 4.**
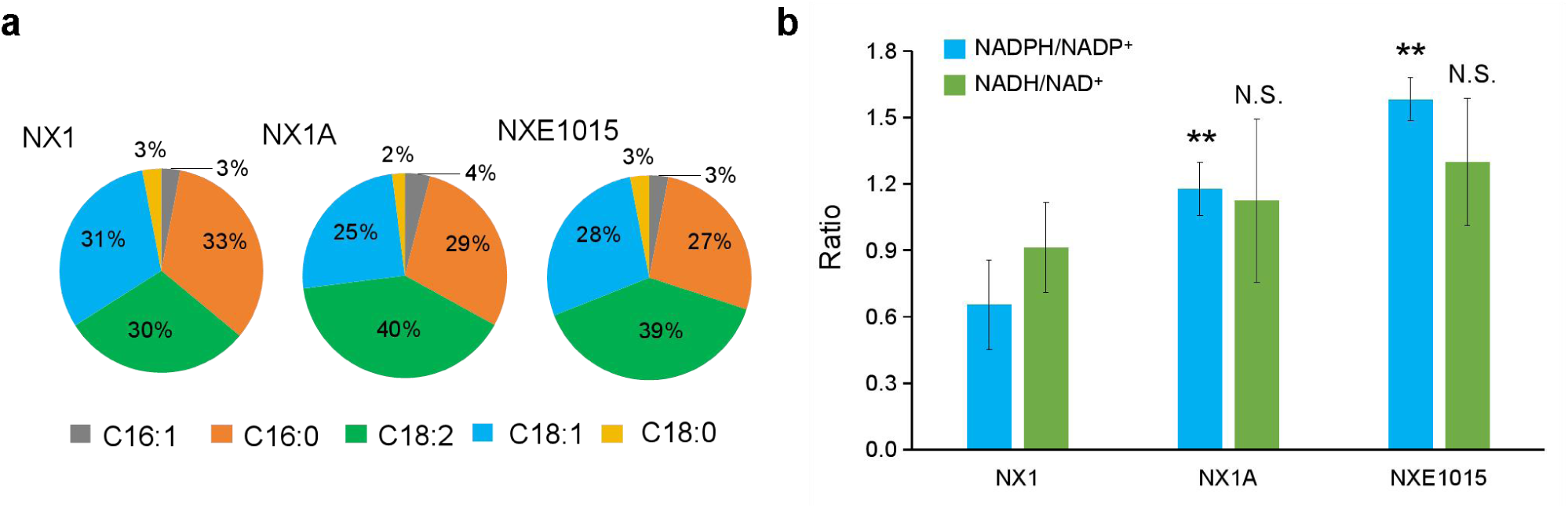
Reverse engineered strain NX1A and the evolved strain NXE1015 exhibit similar trends in terms of FFA production and cofactor levels. (a) The composition of FFAs produced by strains NX1, NX1A and NXE1015. (b) Cofactor measurements of the strains NX1, NX1A and NXE1015. The data are presented as the means ± s.e.ms. (n = 3 biologically independent samples).

**Extended Data Fig. 5.**
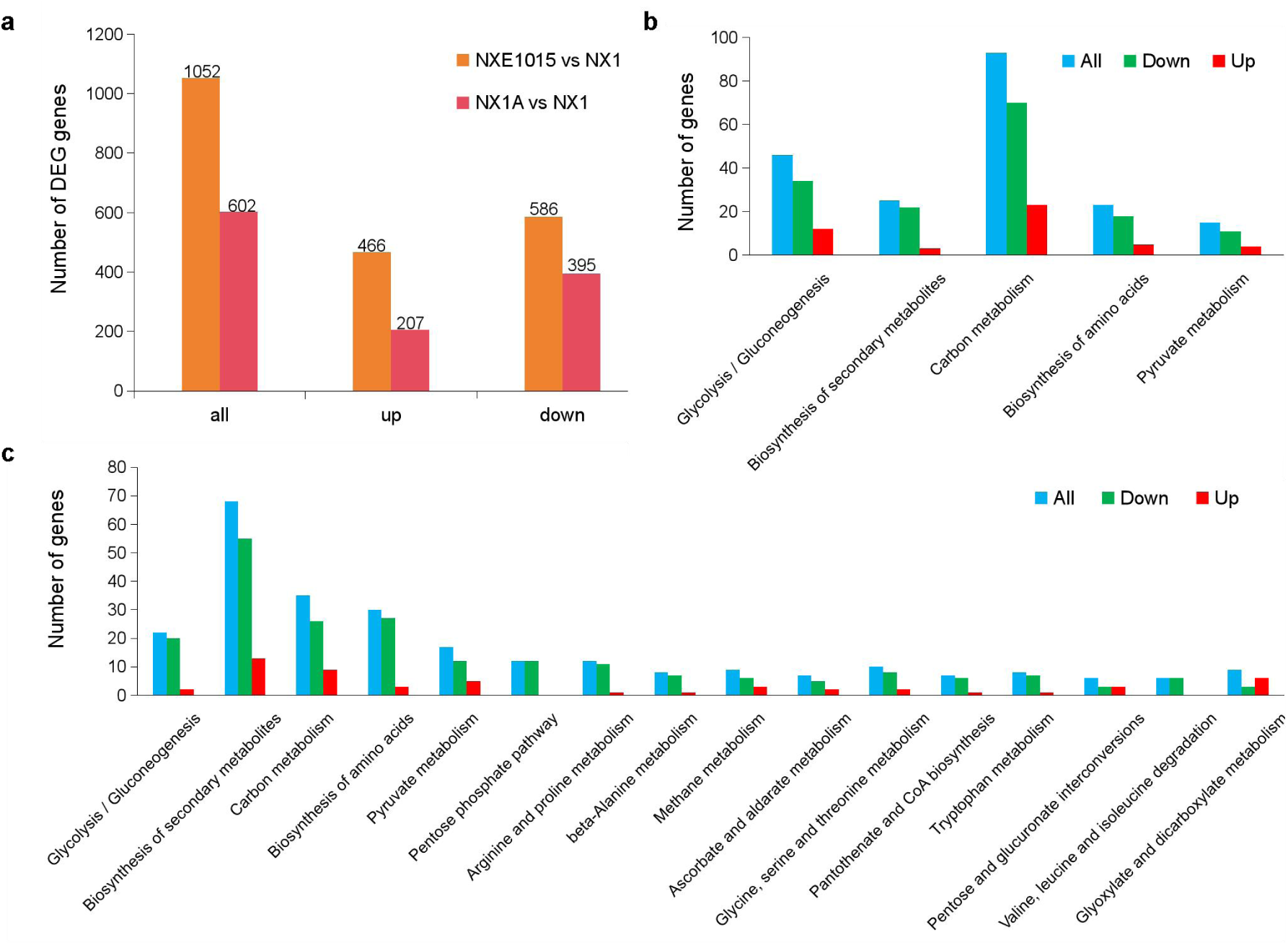
Transcriptome sequencing analysis of differentially expressed genes. (a) Column chart showing the number of differentially expressed genes (DEGs) in strains NXE1015 and NX1A. The number of DEGs in strain NXE1015 was greater than that in NX1A, and more than half of the DEGs were downregulated in both strains NXE1015 and NX1A. DEGs in strains NXE1015 (b) and NX1A (c) were annotated by KEGG pathway analysis. The DEGs in strain NX1A were similar to those in NXE1015, especially those in pathways related to carbohydrate metabolism and the biosynthesis of amino acids. Notably, the KEGG-annotated DEGs in strain NX1A also included more pathways, such as CoA biosynthesis and amino acid metabolism (statistical analysis was conducted by two paired t tests, *p < 0.05, **p < 0.01). The data are presented as the means ± s.e.ms. (n = 3 biologically independent samples).

**Extended Data Fig. 6.**
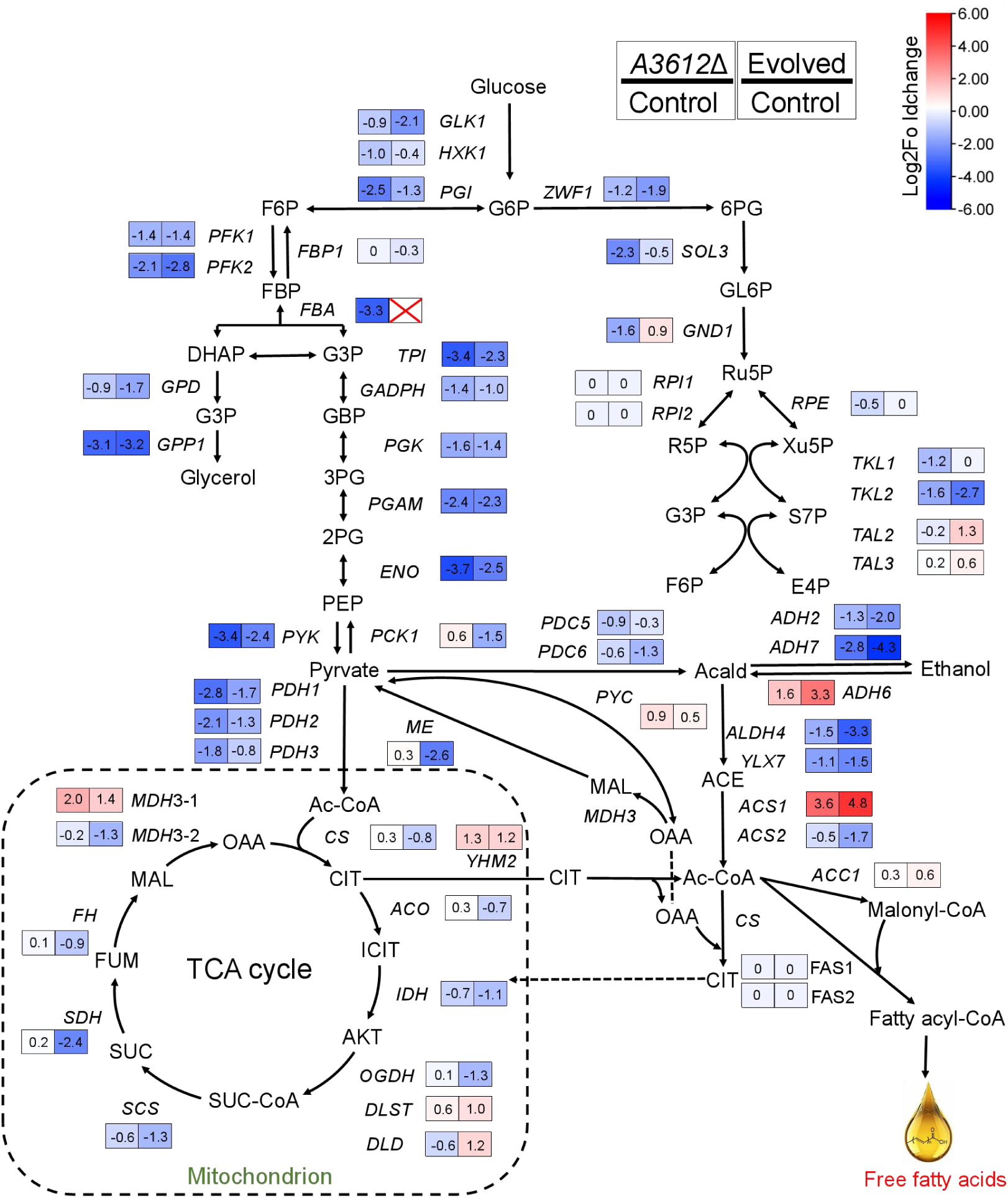
DEGs related to central metabolism and their effects on glucose utilization and FFA production. The results from the transcriptional analysis are shown as relative expression levels in NX1A (reverse engineered strain, Δ*CMR*) or NXE1015 (evolved strain) compared with NX1 (control strain). Gene expression is presented as the log_2_-fold change (p-adj <0.05). The log_2_-fold change (down/up) of the genes is indicated by blue (down) and red (up). The data are presented as the means ± s.e.ms. (n = 3 biologically independent samples).

**Extended Data Fig. 7.**
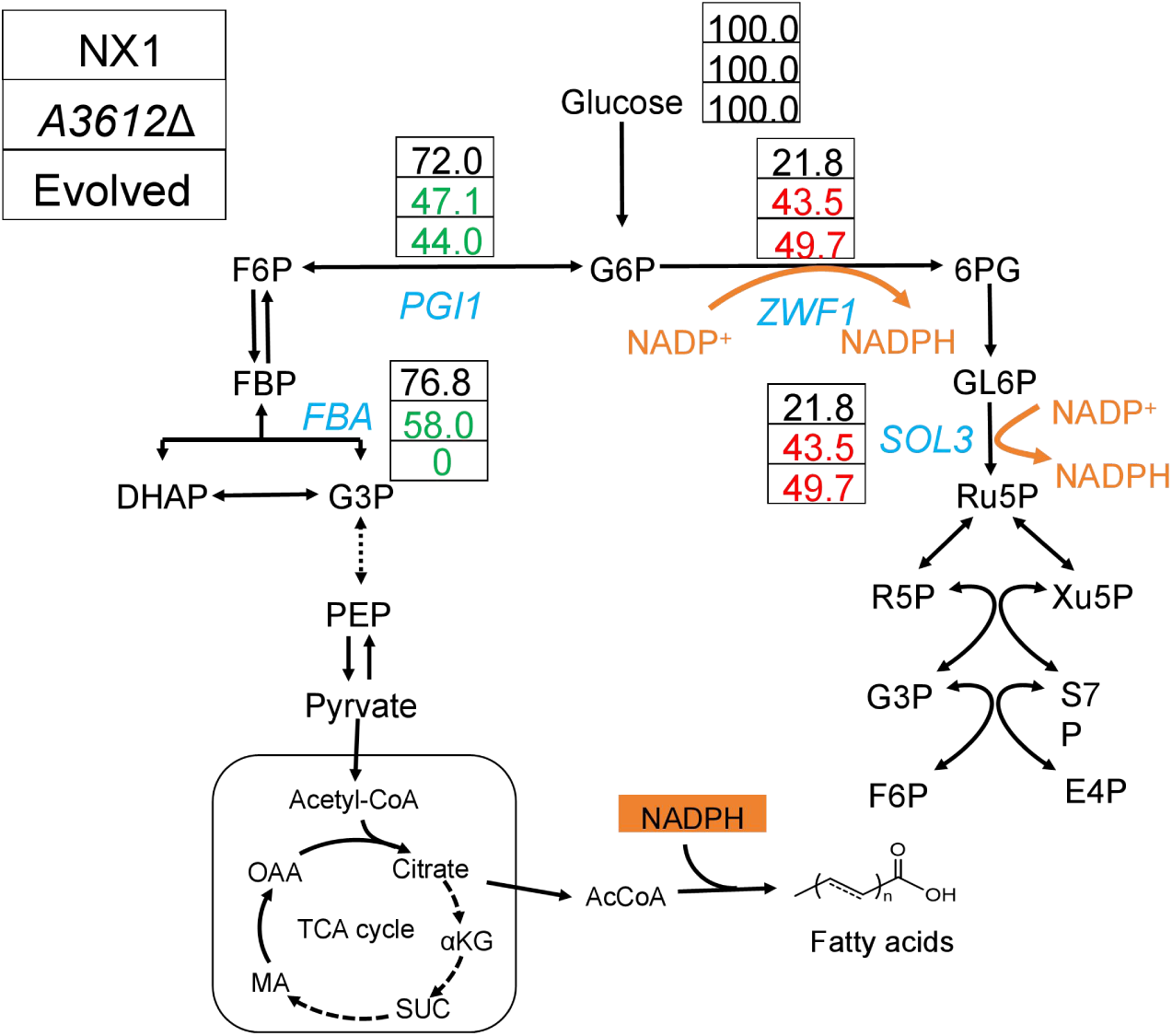
Metabolic flux analysis (MFA) of strains NX1 (control), NX1A (*A3612*Δ) and NXE1015 (evolved). In MFA, the fluxes to the different products were represented relative to the uptake of 100 mmol of glucose. Important pathway genes are highlighted in blue. Red represents an increase in pathway metabolic flux, whereas green represents a decrease in pathway metabolic flux.

**Extended Data Fig. 8.**
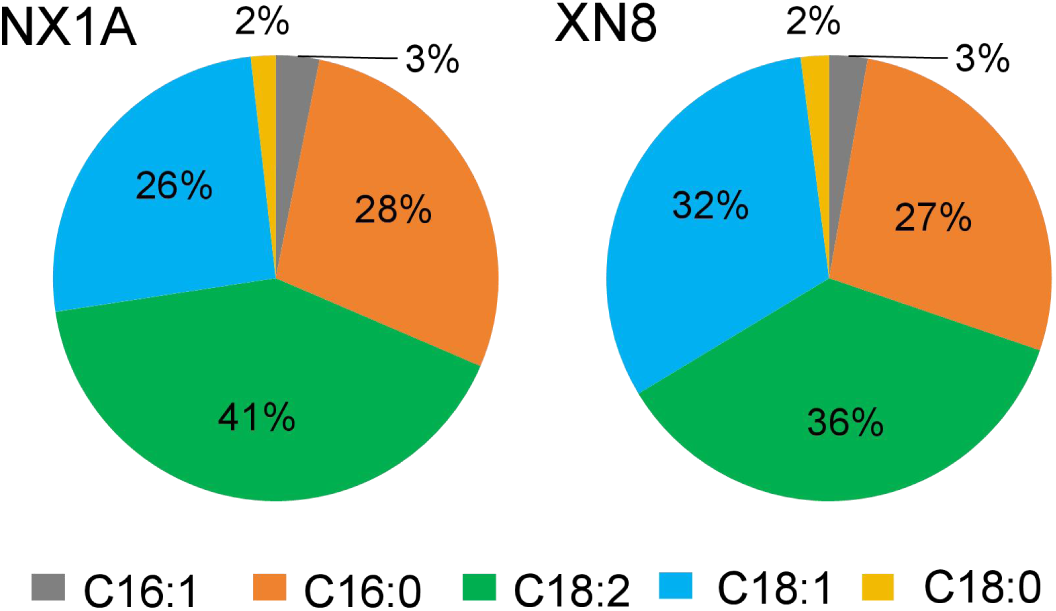
The composition of FFAs in strains NX1A and XN8.

**Extended Data Fig. 9.**
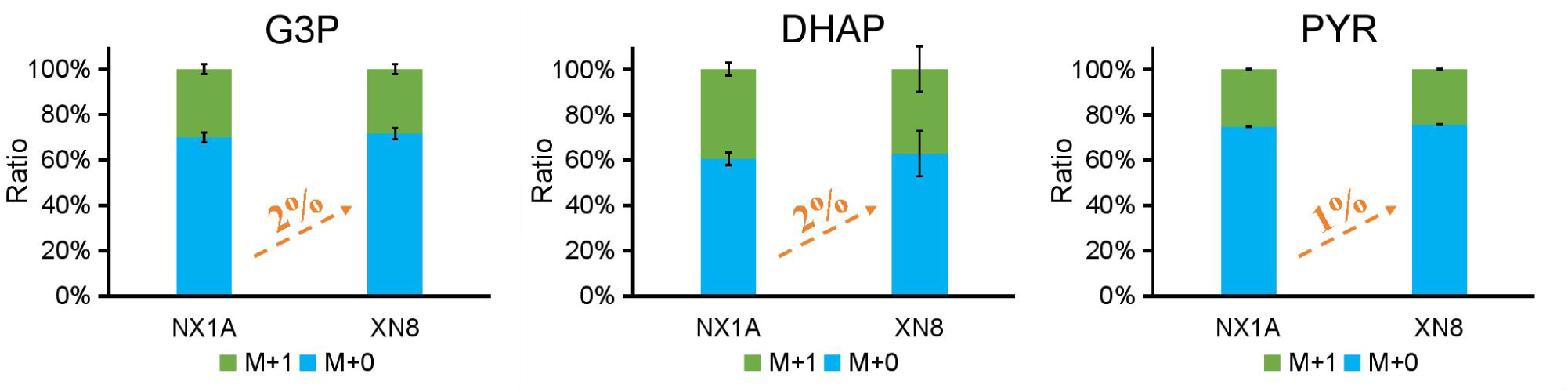
^13^C-labeled glucose tracers. The pathway switching in the strains NX1A and XN8 (P*_GAP_*-*BbXFPK*-*CkPTA*) were validated by isotope tracing experiment. Data are presented as mean ± s.e.m. (n = 3 biologically independent samples).

